# Combining amoxicillin and relebactam provides a new therapeutic option for *Mycobacterium abscessus* infection

**DOI:** 10.1101/563189

**Authors:** Rose C. Lopeman, James Harrison, Savannah E. R. Gibson, Daniel L. Rathbone, Maya Desai, Peter Lambert, Jonathan A. G. Cox

## Abstract

Infections caused by the opportunistic pathogen *Mycobacterium abscessus* are increasing in prevalence within the cystic fibrosis population. Our limited understanding of this ubiquitous environmental microorganism matched with its intrinsic resistance to most classes of antibiotic, including *β*-lactams, has left us with an urgent demand for new, more effective therapeutic interventions. *De novo* antimicrobial drug discovery is a lengthy process and so we have taken the approach of repurposing known antibiotics in order to provide a rapidly implementable solution to a current problem. Here we report a significant potential step forward in the treatment of *M. abscessus* infection by sensitising the organism to the broad spectrum μ-lactam antibiotic, amoxicillin, using the competitive μ-lactamase inhibitor, relebactam. We demonstrate by disk diffusion and broth microdilution assay that this combination works synergistically to inhibit *M. abscessus*. We also demonstrate the direct competitive inhibition of the M. abscessus *β*-lactamase, Bla_Mab_, using a novel thin-layer chromatography-based assay for *β*-lactamase inhibition, which is subsequently validated kinetically by spectrophotometric assay using the nitrocefin reporter assay and *in silico* binding studies. Furthermore, we reverse the sensitivity by overexpressing Bla_Mab_ in *M. abscessus*, demonstrating relebactam-Bla_Mab_ target engagement. Finally, we demonstrate the *in vitro* efficacy of this combination against a collection of *M. abscessus* clinical isolates, as well as the *Mycobacterium tuberculosis* model organism *Mycobacterium bovis* BCG, demonstrating the significant therapeutic potential of the amoxicillin and relebactam combination.

## Introduction

The incidence of infections with the rapidly growing non-tuberculous mycobacterial (NTM) organism *Mycobacterium abscessus* is increasing due to the increase of immunocompromised individuals (Petrini, et al. 2006) (Griffith, et al. 2014). *M. abscessus* is widely responsible for opportunistic pulmonary infections in patients with structural lung disorders such as cystic fibrosis (CF) and bronchiectasis (Griffith, et al., 2007) as well as skin and soft tissue infections (SSTIs) in humans (Moore, 1953) (Fitzgerald, et al., 1995). The ubiquitous environmental nature of M. abscessus may go some way to explaining the high levels of intrinsic drug resistance to most major classes of antibiotic that is observed clinically (Nessar, et al. 2012). Recently, macrolides were added to the ever growing list of ineffective antimicrobial agents for *M. abscessus* infection, exemplifying the urgent need for new ways to treat this infection (Nessar, et al. 2012). Additionally, *M. abscessus* is resistant to two of the frontline antibiotics used for tuberculosis treatment; rifampicin and ethambutol (Nessar, et al. 2012), and many of the new drugs being discovered for treating tuberculosis do not exhibit any antimicrobial activity against *M. abscessus* (Soni, et al. 2016).

Pulmonary *M. abscessus* disease was historically treated with macrolide antibiotics such as clarithromycin (Griffith, et al., 2007), however the steady emergence of antibiotic resistant strains has prompted the implementation of combination therapy using azithromycin (macrolide) along with amikacin (aminoglycoside) and at least one other drug of a different class (Leung, et al. 2013). The British Thoracic Society recommends a biphasic treatment for pulmonary *M. abscessus* infection comprising of a 1-month initiation phase including intravenously administered amikacin, tigecycline and imipenem, supplemented with oral clarithromycin or azithromycin (depending on sensitivity results). This is followed by a 12-month continuation phase comprising of nebulised amikacin, and a combination of 1-4 of clofazimine, linezolid, minocycline, moxifloxacin or co-trimoxaole (Lopeman et al. 2019) (Haworth et al 2017).

The advancements in whole genome sequencing technology have enabled the identification of three distinct subspecies within the *M. abscessus* complex, comprising *M. abscessus* subsp. abscessus, *M. abscessus* subsp. bolletii, and *M. abscessus* subsp. massiliense, the latter of which contains a non-functional erm(41) gene, resulting in a macrolide sensitive phenotype (Bryant et al. 2013) (Tortoli et al. 2016). Mutations in rrs and rrl 16S rRNA genes have been identified corresponding to high levels of aminoglycoside resistance, prohibiting the effective use of amikacin in treatment for *M. abscessus* subsp. abscessus and *M. abscessus* subsp. massiliense clinical isolates specifically (Kehrmann et al. 2016). All *M. abscessus* subsp. contain an endogenous class A *β*-lactamase (Bla_Mab_) conveying intrinsic resistance to the *β*-lactam antibiotics (Soroka et al 2014). The identification of Bla_Mab_, a homolog of the *Mycobacterium tuberculosis* endogenous *β*-lactamase, BlaC, sparked widespread investigation into the potential of *β*-lactamase inhibitors to supplement the treatment of *M. abscessus* infection. A study demonstrated the inhibitory activity of avibactam, a non-*β*-lactam *β*-lactamase inhibitor, against Bla_Mab_ significantly improving the in vitro and in vivo activity of imipenem (Lefebvre et al. 2016). Recently, multiple studies have been published demonstrating a number of *β*-lactam combinations displaying high levels of *in vitro* synergistic activity against *M. abscessus* complex (Pandey et al 2019) (Story-Roller et al. 2019). Furthermore, one of these studies identified the *in vitro* activity of two new non-*β*-lactam *β*-lactamase inhibitors, namely relebactam and vaborbactam, demonstrating synergistic activity between imipenem and relebactam as well as meropenem and vaborbactam (Kaushik et al. 2019).

In this study, we have built on previous work (Kaushik et al. 2019) and identified a significant step forward in treatment potential for *M. abscessus* infection. We demonstrate the *in vitro* capacity for sensitising the organism to the broad spectrum *β*-lactam antibiotic, amoxicillin, using the *β*-lactamase inhibitor, relebactam. We establish synergy between amoxicillin and relebactam as well as activity against a range of *M. abscessus* clinical isolates. We have identified the mechanism of relebactam-Bla_Mab_ inhibition as competitive using recombinant protein in both a novel thin-layer chromatography (TLC)-based assay, as well as previously established spectrophotometric methods. Furthermore, this has been delineated using in silico molecular docking. This offers a considerable therapeutic potential as imipenem, a mainstay of current *M. abscessus* treatment, is currently undergoing phase III clinical trials in combination with relebactam, suggesting the simple addition of amoxicillin and relebactam to the treatment regime has the potential to significantly improve the treatment outcome.

## Materials and Methods

### Bacterial isolates

A total of 16 *M. abscessus* clinical isolates from Brighton and Sussex Medical School and *M. abscessus* NCTC 13031 were used in this study, along with *Mycobacterium bovis* BCG Pasteur. Stock solutions of the isolates were kept in 50% glycerol (Sigma, Dorset, UK) and Middlebrook 7H9 Broth and stored at −80 °C. Isolates were grown in Middlebrook 7H9 medium supplemented with 10% oleic acid-dextrose-catalase (OADC), 1% glycerol (50% w/v) and 0.05% Tween 80 (v/v) prior to testing. *Escherichia coli* Top 10 cells were used for propagation of plasmid DNA. These cells were routinely grown in nutrient broth, or nutrient agar (Oxoid, UK) at 37 °*C. E. coli* BL21 (DE3) cells were used for the overproduction of recombinant protein, grown in Terrific Broth (Melford, UK) at 37 °C.

### Antimicrobials

The antimicrobial agents meropenem (MEM), amoxicillin (AMX) and phenoxymethylpenicillin (Penicillin V/PenV) were obtained from Sigma Aldrich (Dorset, UK) and relebactam (REL) was obtained from Carbosynth (Compton, UK). Stock solutions were prepared in sterile de-ionised water and stored at −20 °C until use.

### Disk diffusions

For the disk diffusion assays, *M. abscessus* clinical isolates were grown in Middlebrook 7H9 Broth to logarithmic phase and 100 μL of bacterial culture was inoculated into 10 mL Middlebrook 7H9 Broth supplemented with 0.7% bacteriological agar. This was poured as a layer in agar plates on top of 15 mL Middlebrook 7H11 Agar supplemented with 10% OADC and 1% glycerol. 6 mm sterile filter paper diffusion disks were placed on the agar and were subsequently impregnated with antibiotic at the following concentrations in sterile distilled water: relebactam 1 μL of 10 mg/mL, amoxicillin 3.3 μL of 3 mg/mL and meropenem 1 μL of 10 mg/mL. Plates were incubated at 30 °C for 5 days or until clear zones of inhibition were visualised. Zones of inhibition were measured across the diameter to include the disk itself. Disk diffusions were also performed on M. abscessus pVV16-*bla*_*Mab*_ and *M. abscessus* pVV16.

### Broth microdilution assay

The broth microdilution assay was performed as described previously with alterations appropriate to this study (Institute, Clinican and Laboratory Standards 2012) (Caleffi-Ferracioli et al. 2013). Plates were prepared by serially diluting amoxicillin (0.128 mg/mL – 0 mg/mL) or meropenem (0.128 mg/mL – 0 mg/mL) in the x-axis and relebactam (0.128 mg/mL – 0 mg/mL) in the y-axis. 80 μL of Mycobacterium abscessus NCTC 13031 or Mycobacterium bovis BCG suspension adjusted to an OD_600_ of 0.1-0.2 was inoculated in each well to reach a final well volume of 100 μL. Plates were sealed and incubated at 30 °C for 5 days for *M. abscessus* cultures or 37 ^°^C for 10 days for M. bovis BCG. The MIC of amoxicillin/relebactam combination was determined by optical density measurement using a spectrophotometric plate reader and identified as the well that had the lowest concentrations of both compounds and exhibited no bacterial growth. The relevant well was retroactively plotted into a growth curve over time, and this growth curve was compared to the wells containing no drug, and the wells containing relebactam only and amoxicillin only. The broth microdilution assay was also performed on *M. abscessus* pVV16-*bla*_Mab_ and *M. abscessus* pVV16.

### Generation of *M. abscessus* pVV16-*bla*_*Mab*_ overexpressor strain

The *bla*_*Mab*_ gene (MAB_2875) from *Mycobacterium abscessus* NCTC 13031 was amplified by polymerase chain reaction (PCR). Primers used were as follows (with restriction site underlined): Forward primer AAAAAA<u>GGATCC</u>GCGCCGGACGAACTCGCC and Reverse primer AAAAAA<u>AAGCTT</u>AGCGCCGAAGGCCCGCAG (Eurofins Genomics). Amplicons were purified and cloned into pVV16 using *Bam*HI/*HinDIII* restriction sites and the sequence was confirmed (Eurofins Genomics). Both the plasmid pVV16 and the construct pVV16-*bla*_*Mab*_ were inserted into *M. abscessus* NCTC 13031 cells by electroporation (2.5kV, 2.5 μF and 1000 Ω).

### Expression and purification of recombinant Bla_Mab_

The *bla*_*Mab*_ gene, with the first 90 base pairs omitted (resulting in a −30 residue N-terminal truncated protein), was amplified by PCR using the following primers (with restriction sites underlined): Forward primer AAAAAA<u>GGATCC</u>GCGCCGGACGAACTCGCC and Reverse primer AAAAAA<u>AAGCTT</u>TCAAGCGCCGAAGGCCCG (Eurofins Genomics). Amplicons were purified and cloned into pET28a using *Bam*HI/*HinDIII* restriction sites and the sequence was confirmed (Eurofins Genomics). Bla_Mab_ was expressed in *E. coli* BL21 (DE3) cells by addition of 1 mM Isopropyl β-D-1-thiogalactopyranoside (IPTG) and incubation at 25 °C for 18 h. Bla_Mab_ was purified by Immobilised Metal Affinity Chromatography (IMAC) and dialysed into 25 mM Tris HCl pH 7, 100 mM NaCl.

### Thin Layer Chromatography (TLC) activity assay

Relebactam (2 mg/mL) was added to recombinant Bla_Mab_ (0.01 mg/mL) and incubated for 5 minutes at room temperature, before addition of Penicillin V (4 mg/mL) for a further 10 minute incubation at room temperature. Alongside appropriate control reactions (Figure 2), 1 μL of the reaction was spotted onto aluminium backed silica gel plates (5735 silica gel 60 F_254_, Merck) and dried before being subjected to Thin Layer Chromatography (TLC) using ethyl acetate:water:acetic acid (C_4_H_8_O_2_:H_2_O:CH_3_COOH) (3:1:1, v/v/v). Once dry, plates were visualised by being dipped into KMnO_4_ TLC stain with light charring.

### Biochemical analysis of Bla_Mab_ inhibition by Relebactam

Recombinant Bla_Mab_ (0.25 nM) was mixed with an increasing concentration of relebactam (0, 0.1, 1, 10 and 100 μM) and 100 μM nitrocefin (Carbosynth, UK). The hydrolysis of nitrocefin was monitored at 486 nm using a Multiskan Go plate reader (Thermo Scientific). This was repeated using a varying concentration of nitrocefin (1-500 μM) with a shorter range of relebactam concentrations (0, 0.5, 0.75, 1 and 2.5 μM) and initial velocities (v_i_) were plotted against substrate concentration. Data analysis was performed as previously described (Dubée et al., 2014, Papp-wallace et al., 2018). Data analysis was conducted using Graphpad Prism 7.

The decarbamylation rate was determined by incubation of Bla_Mab_ (1 μM) with relebactam (20 μM) for 1 hour at 25 °C. The reaction mixture was then subsequently diluted both 10,000 and 50,000 fold (1.8 nM and 0.36 nM relebactam with 90 pM and 18 pM Bla_Mab_ respectively), before addition of 100 μM nitrocefin. The reaction was monitored at 486 nm using a Multiskan Go plate reader (Thermo Scientific).

### *In Silico* Modelling of the Possible Interaction of Relebactam with the Bla_Mab_

The Bla_Mab_ X-ray crystal structure was obtained from the Protein Data Bank, accession code 4YFM. The A chain was submitted to the GHECOM pocket-finding server (GHECOM 1.0) (Kawabata 2010; Kawabata and Go 2007). The top six pockets identified were used to define the target site for protein-ligand docking experiments between the enzyme and relebactam using CACHe Worksystem Pro (version 7.5.0.85, Fujitsu Ltd). The amino acid residues lining theses pockets are given in Figure 3(c). Pocket 1 corresponded to the main (catalytic) site in the enzyme and for the purposes of the docking experiment was redefined as all amino acid residues within 8 Å of serine 71. Hydrogen atoms were added using the default settings in line with presumed protonation states for ionisable amino acid side-chains. The positions of the added hydrogen atoms were optimised by locking the coordinates of all the non-hydrogen atoms and subjecting the system to a molecular mechanics (MM2) geometry optimisation. Relebactam was docked four times into each pocket, using CACHe Worksystem Pro. The amino acid side-chains in each pocket were allowed to be flexible as were all rotatable bonds in relebactam. The genetic algorithm settings for the docking protocol included population size 50, maximum generations 3000, crossover rate 0.8, mutation Rate 0.2 and convergence when the RMSD population fitness was less than 1. The best-scoring consensus complexes from each series of dockings were taken forward for molecular dynamics simulation. The required input files were prepared using the Antechamber module of the AMBER Tools package (Version 14; Case et al, 2005), implementing the ff14SB force field. The system was neutralised by addition of sodium ions and then solvated within a truncated octahedron of TIP3P water molecules extending 8 Å from the surface of the protein. Using the Amber 14 molecular dynamics package CUDA version (Salomon-Ferrer et al, 2013; Goetz et al, 2012; Le Grand et al, 2013), the system was energy-minimised for 2,000 cycles using a non-bonded cut-off of 12 Å and then heated under constant volume to 300 K over 25 ps under Langevin dynamics (time step = 1 fs). The heating was continued at 300 K for at least 200 ns under constant pressure also using Langevin dynamics (SHAKE on, time step = 2 fs) using the Particle–Mesh–Ewald (PME) method to treat the long range electrostatic interactions with a 12 Å non-bonded cut-off.

## Results

In this paper we have identified that *M. abscessus* can be sensitised to amoxicillin by the addition of the competitive *β*-lactamase inhibitor, relebactam. Furthermore we have shown that meropenem sensitivity can be enhanced by the addition of relebactam. Firstly, we conducted disk diffusion assays with amoxicillin and meropenem with and without relebactam. Subsequently the zones of inhibition (ZOI) were measured as a marker of sensitivity. Amoxicillin alone failed to demonstrate any kind of sensitivity, however the addition of relebactam provided a clear ZOI, demonstrating the induced sensitivity to amoxicillin by relebactam. A ZOI was visible for meropenem, however with the incorporation of relebactam, this ZOI was significantly enhanced, suggesting increased sensitivity upon combination with relebactam (1a). This led us to the hypothesis that relebactam was inhibiting the *M. abscessus β*-lactamase, Bla_Mab_, preventing hydrolysis of amoxicillin to the inactive amoxicilloic acid (1b). In order to assess the clinical relevance of these observations, both amoxicillin and meropenem were tested alone and in combination with relebactam against a panel of clinical isolates of *M. abscessus* obtained from patients at Brighton and Sussex Medical School in addition to *M. abscessus* NCTC strain. In all cases, the addition of relebactam sensitised the isolate to amoxicillin (*n*=3) (1c left). The same experiment was conducted with meropenem, and in most cases, an increase in sensitivity was observed (*n*=3) (1c right). All necessary controls were conducted and no ZOI was observed for relebactam alone.

In order to further validate our observation, liquid cultures were exposed to increasing concentrations of amoxicillin and meropenem with and without a dose response of relebactam (1d top and bottom respectively). As a control, relebactam was also tested for inhibitory activity on its own. These cultures were read every 24 hours using a spectrophotometer at OD_600nm_ up to 96 hours. Relebactam at 128 μg/mL demonstrated no inhibitory activity, giving a growth profile much the same as the compound-free control (no drug), confirming that relebactam lacks antibacterial activity against *M. abscessus*. 32 μg/mL of amoxicillin appeared to show a moderate enhancement of bacterial growth, however when combined with 2 μg/mL of relebactam, this amoxicillin concentration was found to display potent antibacterial activity against *M. abscessus*. The same experiment was conducted with meropenem, wherein partial growth impairment was observed at 8 μg/mL, however when combined with 2 μg/mL relebactam, significant antibacterial activity was observed at this concentration. End point statistical analysis was conducted using a t-test to assess the significance of differences between cultures with and without relebactam. In both cases, a statistically significant increase in sensitivity was observed upon addition of 2 μg/mL of relebactam in combination with amoxicillin or meropenem. This experiment was repeated with a Bla_Mab_ overexpressing strain to prove direct binding of relebactam to Bla_Mab_ (1e). The *M. abscessus* containing the Bla_Mab_ constitutive overexpression plasmid pVV16-*bla*_*Mab*_ lost sensitivity to the amoxicillin-relebactam combination compared to the empty vector control strain (pVV16-). This result is replicated using the ZOI method (1c), clearly demonstrated drug-target interaction between relebactam and Bla_Mab_. Plate images and plate map for the subsequent resistance induced by Bla_Mab_ overexpression are shown in Supplementary Figure 1. To briefly explore the wider application for the amoxicillin-relebactam combination against other mycobacterial infections, we tested its activity in liquid culture against the widely used model organism for Mycobacterium tuberculosis, *Mycobacterium bovis* BCG (Supplementary Figure 3). Our results show that *M. bovis* BCG amoxicillin sensitivity is enhanced from 200 to 25 μg/mL upon addition of 8 μg/mL relebactam.

**Figure 1:**
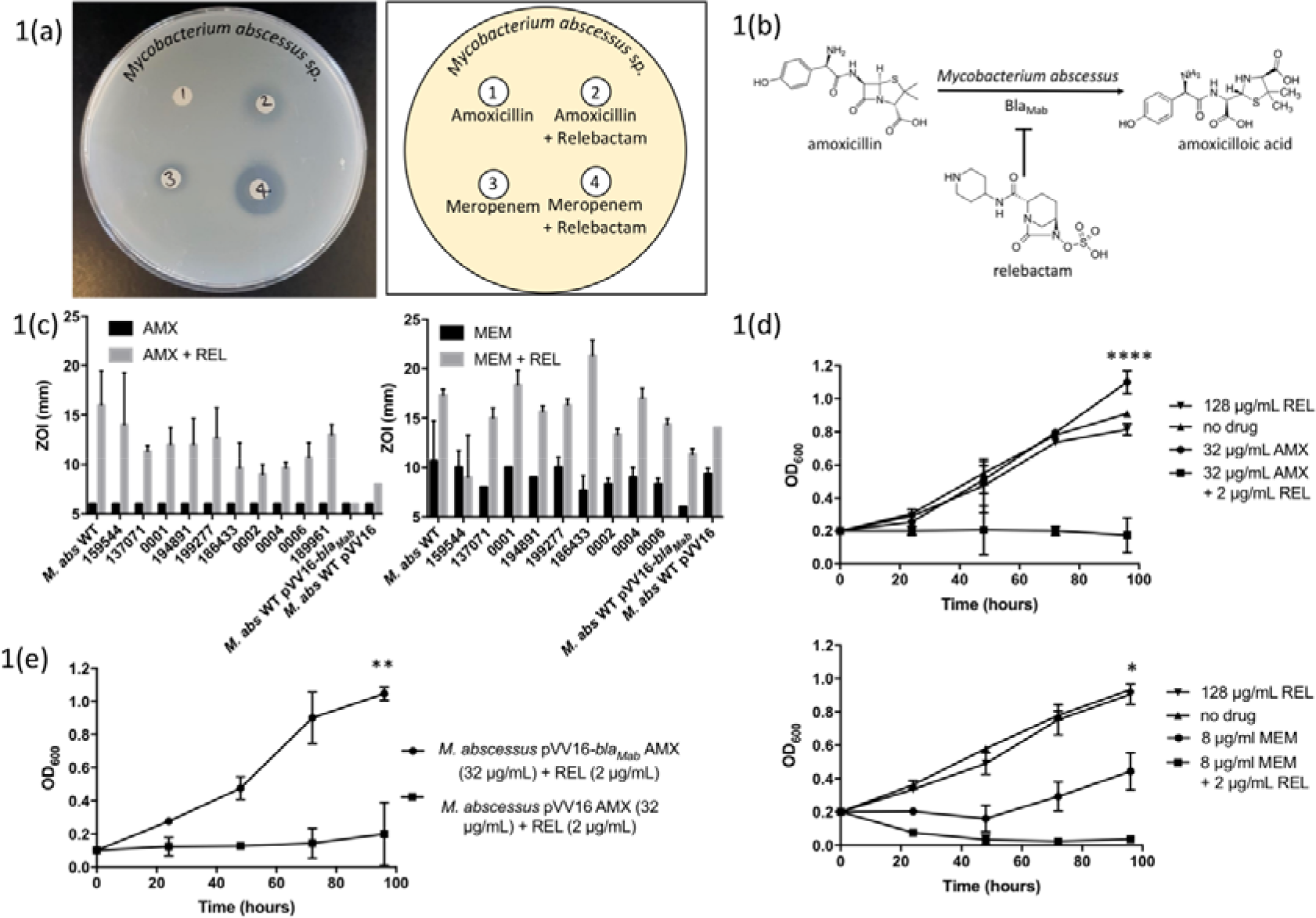
Relebactam sensitises *Mycobacterium abscessus* to amoxicillin and improves sensitivity to meropenem. A disk diffusion experiment (1a) and corresponding plate map demonstrating enhanced sensitivity of *M. abscessus* by zone of inhibition to amoxicillin (1) with the addition of relebactam (2) and meropenem (3) with the addition of relebactam (4). The inhibition of *M. abscessus* -lactamase Bla_Mab_ by relebactam is summarised (1b). The disk diffusion experiments were conducted with the NCTC *M. abscessus* strain along with a panel of clinical isolates and *M. abscessus* pVV16-*bla*_Mab_ and *M. abscessus* pVV16. The diameter of the zones of inhibition (ZOI) measured in millimetres. All strains of *M. abscessus* tested, except *M. abscessus* pVV16-*bla*_*Mab*_, were resistant to amoxicillin (AMX) alone, however displayed sensitivity with the addition of relebactam (REL). *M. abscessus* pVV16-*bla*_*Mab*_ was resistant to amoxicillin both with and without relebactam (1c). The same was conducted for meropenem (MEM) and likewise, activity was clearly enhanced with the addition of relebactam, even in the case of *M. abscessus* pVV16-*bla*_*Mab*_ which was resistant to meropenem alone but sensitised with the addition of relebactam (1c). Growth curves were conducted with *M. abscessus* NCTC in medium containing 128 relebactam and 32 amoxicillin with and without relebactam at 2 μg/mL. Growth inhibition was only observed with relebactam in combination with amoxicillin (1d). Likewise, the inhibitory activity of meropenem was clearly enhanced with the addition of relebactam (1d). A t-test was used for end point analysis between samples +/− relebactam and the results were deemed to be significant with *p* values of <0.0001 and 0.0152 for amoxicillin and meropenem respectively. The same growth curves were performed with *M. abscessus* pVV16-*bla*_*Mab*_ and *M. abscessus* pVV16 in medium containing 32 μg/mL amoxicillin with and without relebactam at 2 μg/mL (1e). *M. abscessus* pVV16-*bla*_Mab_ displayed resistance to amoxicillin and relebactam together whereas *M. abscessus* pVV16 retained sensitivity to amoxicillin and relebactam (*p* = 0.0017).

**Figure 2.**
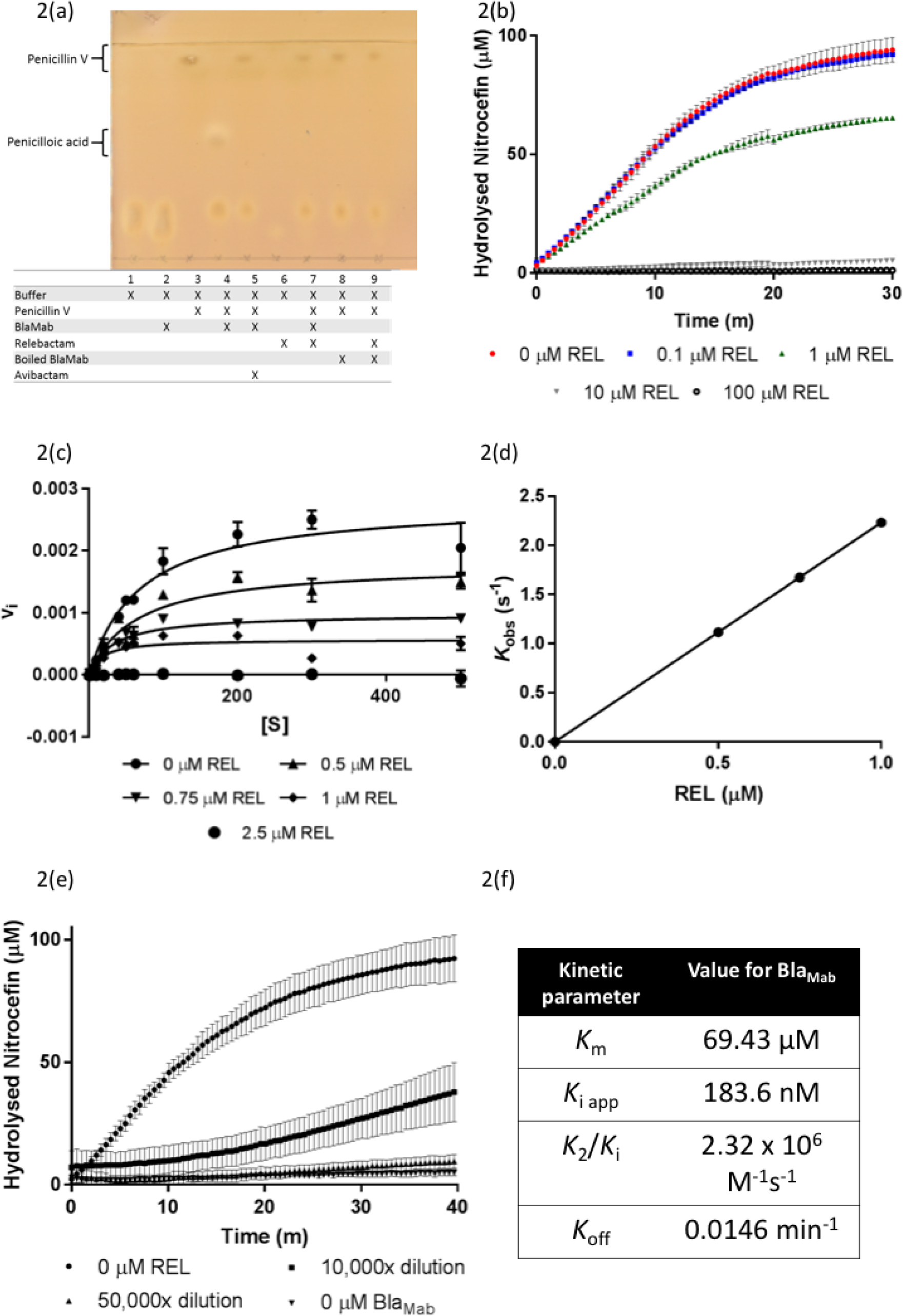
Biochemical analysis of relebactam inhibition of *M. abscessus β*-lactamase, Bla_Mab_. Our novel Thin Layer Chromatography (TLC) assay (2a) exhibiting the activity of Bla_Mab_ in the turnover of penicillin V (high R_f_ value) to penicilloic acid (lower R_f_ value). In the absence, or termination of activity of Bla_Mab_ (by boiling (100 °C) for 1 h) or addition of known inhibitor avibactam (Lefebvre et al., 2017) (200 μg/mL) no lower R_f_ value spot corresponding to penicilloic acid is seen on the TLC plate. The addition of relebactam to the reaction between Bla_Mab_ and penicillin V also results in the absence of the lower Rf value spot, suggesting inhibition of Bla_Mab_ (2a). This observed inhibition is validated by a spectrophotometric analysis, where the turnover of nitrocefin (100 μM) by Bla_Mab_ (0.25 nM) was monitored at 486 nm with a varying concentration of relebactam (0, 0.1, 1, 10 and 100 μM). The increase in concentration of relebactam resulted in partial inhibition of nitrocefin turnover at 1 μM and total abrogation at 10 μM (2b). The initial velocity (v_i_) of the reaction between Bla_Mab_ and nitrocefin was monitored for a range of substrate concentrations (1-500 μM) and relebactam concentrations (0, 0.5, 0.75, 1 and 2.5 μM). This data was plotted v_i_ vs [S] in order to determine *K*_m_ values (2c/f). The values for *K*_obs_ were obtained as previously described (Dubée et al., 2014) and plotted against relebactam concentrations ([I]) to deduce a carbamylation rate (*K*_2_/*K*_i_) for Bla_Mab_ (2d). The kinetics of Bla_Mab_ decarbamylation were assessed to show the recovery of nitrocefin hydrolysis by Bla_Mab_ after inhibition by relebactam in order to derive a *K*_off_ value (2e). Kinetic parameters were derived as described previously (2f) (Dubée et al., 2014).

**Figure 3:**
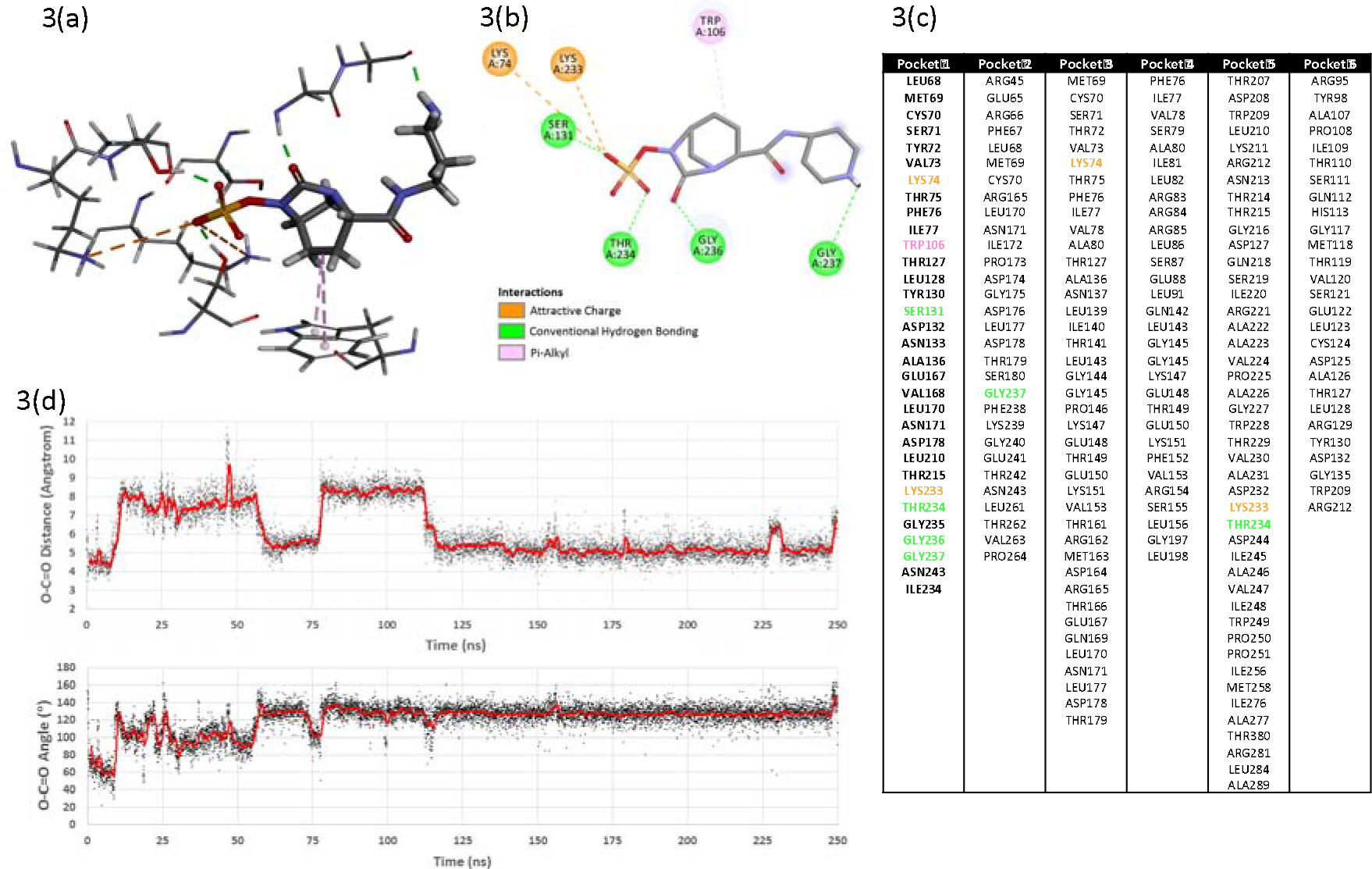
*In silico* modelling of the possible interaction of relebactam with the *M. abscessus* β-lactamase, Bla_Mab_. 3D and 2D protein-ligand interaction diagrams for relebactam in the main (catalytic) active site after molecular dynamics simulation (3a and 3b respectively). Amino acid residues featured in the top six potential binding pockets identified for Bla_Mab_. Pocket 1 corresponded to the main (catalytic site) in the enzyme and for the purposes of the docking experiment was redefined as all amino acid residues within 8 Å of serine 71 (3c). Time-courses of the serine 71 hydroxyl oxygen – relebactam carbonyl carbon distance and the corresponding O-C=O angle. The actual values are plotted as black points and a 50-frame moving average is over-plotted in red (3d).

In order to validate the inhibitory activity observed phenotypically in Figure 1, we conducted biochemical analysis of the activity of relebactam on the *M. abscessus* endogenous *β*-lactamase, Bla_Mab_. The gene was amplified by PCR, digested, ligated into pET28a and sequenced, before transformation into chemically competent *E. coli* BL21. Bla_Mab_ was expressed, cells harvested and the enzyme purified by IMAC. We subsequently devised a novel TLC-based assay for assessing *β*-lactamase activity by separating the penicillin V substrate from the penicilloic acid product. This assay enabled us to demonstrate the *β*-lactamase activity of our purified Bla_Mab_ (2a), as well as assay the efficacy of inhibitors against the enzyme (Supplementary 2a and 2b). We used avibactam as a positive control, as its inhibitory activity against Bla_Mab_ has previously been described by Lefebvre et al. 2017. Lane 1 contained protein purification buffer only, and lane 2 had the addition of enzyme (0.01 mg/mL). Following chromatography, two lower spots are observed in lanes 1 and 2, indicative of buffer. Penicillin V was added to lane 3 and gave a characteristic spot of high R_f_ value, demonstrating unhydrolysed penicillin resulted in a spot just below the solvent front. Lane 4 was identical to lane 3 with the addition of Bla_Mab_ protein. The hydrolysis of penicillin V to penicilloic acid by Bla_Mab_ resulted in a spot with reduced R_f_ value. The pre-incubation of enzyme with avibactam in lane 5 resulted in a loss of the lower penicilloic acid spot, demonstrating the inhibition of Bla_Mab_ activity. In lane 6, relebactam alone did not resolve on the TLC, but its pre-incubation with Bla_Mab_ before addition to the penicillin substrate in lane 7 resulted in total inhibition of hydrolysis as observed with avibactam (lane 5). Finally, the inhibition of Bla_Mab_ is further validated by the repeat of lane 4 and lane 7 conditions with heat-denatured Bla_Mab_ (lanes 8 and 9 respectively). This result confirms the direct inhibition of Bla_Mab_ by relebactam (*n*=5).

We investigated the kinetic parameters of the Michaelis-Menten kinetics, the apparent *K*_i_ (*K*_i app_), the second-order carbamylation rate constant (*K*_2_/*K*_i_) and the decarbamylation rate constant (*K*_off_) of relebactam inhibition of soluble recombinant Bla_Mab_ using a commercially available β-lactamase substrate, nitrocefin, as a reporter substrate. Nitrocefin is selectively hydrolysed by β-lactamases, resulting in an increase in absorbance which can be monitored at 486 nm. By pre-incubating Bla_Mab_ enzyme with a dose response of relebactam (from 0-100 μM), before initiation of the absorbance assay with the addition of nitrocefin substrate, we observed partial inhibition at 1 μM (0.348 μg/mL) and a complete loss of activity at 10 μM (3.48 μg/mL), confirming the direct inhibitory activity of relebactam on Bla_Mab_ (2b).

The initial velocities (v_i_) of nitrocefin turnover were recorded for a range of nitrocefin concentrations (1-500 μM) over a range of relebactam concentrations (0-2.5 μM). These results were plotted as classical Michaelis-Menten curves (v_i_ vs [S]) (2c), resulting in Michaelis constants (*K*_m_) between 69.43-16.96 μM, before initial velocities were abolished at 2.5 μM relebactam. However, we were unable to derive a *K*_i_ value using this data. In response, we plotted the reciprocal initial velocities against relebactam concentrations as a linear equation and derived *K*_i app_ observed from the Y intercept/slope, which was then normalised for the use of nitrocefin (2f) (Papp-Wallace et al., 2018). The *K*_i app_ value obtained of 183.6 nM is indicative of the high inhibitory potency of relebactam against Bla_Mab_. The carbamylation rate constant (*K*_2_/*K*_i_) of 2.32 × 10^6^ M^−1^s^−1^ is similar to the observed rate for avibactam inhibition of Bla_Mab_ (4.9 × 10^5^ M^−1^s^−1^) (Dubée et al., 2014). The rate of decarbamylation (*K*_off_) of Bla_Mab_ for relebactam was 0.0146 min^−1^, which is again similar to the rate previously obtained for that of avibactam (0.047 min^−1^) (Dubée et al., 2014). Our kinetics analysis show that, like avibactam, relebactam is a potent, competitive and reversible inhibitor of Bla_Mab_, displaying a reasonably rapid “on” rate and slow “off” rate, with only half the enzyme recovering activity after 40 minutes in the absence of relebactam (2e) (Dubée et al., 2014).

Our TLC-based *β*-lactamase assay enabled us to further explore the parameters of the inhibitory activity of relebactam by varying the time of pre-incubation of relebactam with Bla_Mab_ (Supplementary 2a) and the minimum inhibitory concentration required to abrogate catalytic turnover of the penicillin V substrate to the penicilloic acid substrate (Supplementary 2b). We found that penicillin V turnover was rapid and that only addition of relebactam at the same time as the substrate demonstrated turnover of penicillin V, suggesting competitive, reversible inhibition of Bla_Mab_ by relebactam. The dose response of relebactam demonstrated total inhibition down to 20 μg/mL, and activity of Bla_Mab_ was maintained below a relebactam concentration of 2 μg/mL. This suggested a minimal concentration of relebactam required to inhibit Bla_Mab_ in the assay is within the range of 20 to 2 μg/mL. This corresponds to a less than or equal to 100 fold stoichiometric excess of relebactam required to completely inhibit Bla_Mab_ (0.5 μM BlaMab to 57.5 μM relebactam corresponding to 20 μg/mL).

In order to further investigate the mechanism of relebactam inhibition of Bla_Mab_, we conducted molecular docking simulations *in silico*. 6 potential binding sites were identified (3(c)). For pockets 2-6 the ligand was weakly-held and generally exited the pocket after a few tens of nanoseconds. For pocket 1 (corresponding to the main active site) the ligand reoriented itself relative to the docked conformation and thereafter remained relatively stable within the pocket. The binding interactions for the stable pose are shown in Figure 3(a) and 3(b) and include several polar interactions with the sulphonamide moiety and hydrophobic interactions between the relebactam central piperidine ring and tryptophan 106. In addition, after the initial reorientation, the relebactam carbonyl carbon remained in the vicinity of the hydroxyl oxygen of the catalytically-active serine 71 as can be seen after 120 ns in the distance plot given in Figure 3(d). The average distance in this period was approximately 5.5 Å and there were many close approaches. Furthermore, the corresponding O-C=O angle (Figure 3d) in the same period was roughly 130 ° and as such it is reasonable to assume that it would be possible for the serine hydroxyl to attack the relebactam carbonyl and effect a ring-opening. A further molecular dynamics simulation of the enzyme in a periodic box of explicit water was undertaken but this time in the presence of ten copies of unbound relebactam. Over the course of a 200 ns simulation, one ligand found its way into the main active site (pocket 1) after 130 ns and remained stable therein. The other nine copies of the ligand found no place to reside in the enzyme.

## Discussion

In this paper we have demonstrated the inhibitory activity of the non-*β*-lactam based *β*-lactamase inhibitor, relebactam against *M. abscessus* endogenous *β*-lactamase, Bla_Mab_. Alone, this is not sufficient to kill *M. abscessus*, however its use as part of a combination opens up considerable therapeutic potential. We have identified that a low concentration of relebactam co-administration is capable of sensitising *M. abscessus* NCTC and clinical isolates to amoxicillin, well within the therapeutic range for this versatile and widely available antibiotic. Furthermore, we demonstrate relebactam provides a significant increase in inhibitory activity of meropenem, a mainstay of *M. abscessus* clinical intervention. Our study introduces a completely novel TLC-based *β*-lactamase inhibition assay, validated with the commercially available and widely used nitrocefin assay, to investigate the parameters of relebactam’s inhibitory activity. The results of both the kinetics analysis and the *in silico* binding studies demonstrate the mechanism of inhibition of Bla_Mab_ by relebactam as competitive reversible, displaying a reasonably rapid “on” rate and slow “off” rate, indicative of the high potency of the inhibitor against Bla_Mab_.

Relebactam is currently in phase III clinical trial for administration in combination with imipenem, another mainstay in *M. abscessus* front line chemotherapy. Our findings therefore represent a timely and highly impactful discovery that is easily translatable into the clinical setting, not only providing a new therapeutic option for *M. abscessus* infections, but with the potential to treat other mycobacterial infectious diseases such as tuberculosis.

## Author Contributions

R. C. L., J. H., M. D., D. L. R, P. L. and J. A. G. C. intellectually conceived and designed the experiments. R. C. L., J. H., S. E. R. G, D. L. R. and J. A. G. C. conducted the experiments. R. C. L., J. H., D. L. R. and J. A. G. C. wrote the manuscript.

## Funding

This research was funded by Birmingham Women’s and Children’s Hospital Charity Research Foundation (BWCHCRF) (R. C. L. 50% PhD Studentship, match funded by Aston University Prize Scheme) and the Academy of Medical Sciences and Global Challenges Research Fund with a Springboard Grant (SBF003\1088:).

## Acknowledgements

J. A. G. C. is grateful to the Academy of Medical Sciences, Global Challenges Research Fund and Birmingham Women’s and Children’s Hospital Charity Research Foundation (BWCHCRF) for their continued support of the Mycobacterial Research Group at Aston University. The *Mycobacterium abscessus* clinical isolates used in this study were provided by Dr Simon Waddell, Brighton and Sussex Medical School. We are also grateful to Dr Amit Chattopadhyay, Aston University, for technical support.

