## Supplementary Data for "Combining amoxicillin and relebactam provides a new therapeutic option for *Mycobacterium abscessus* infection"

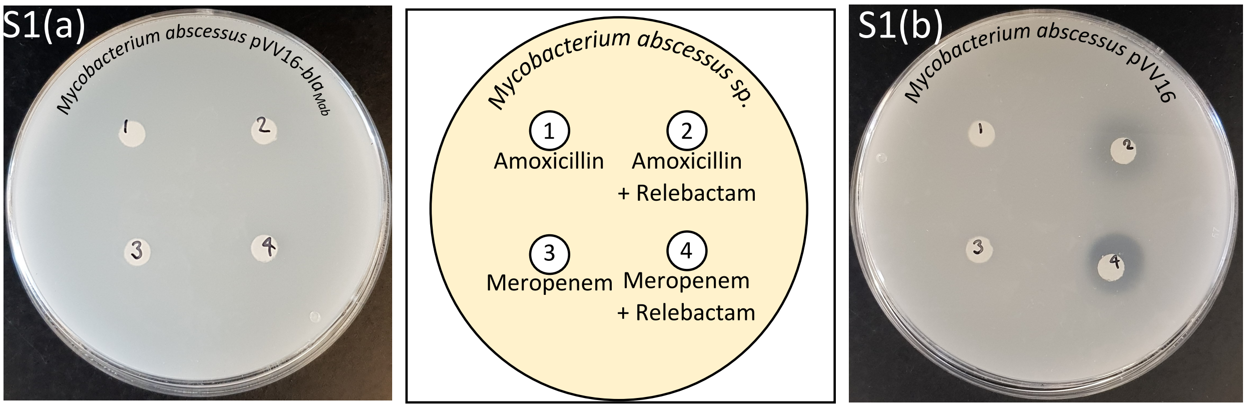


**Supplementary Figure 1: Overexpression of *M. abscessus* β-lactamase results in loss of relebactam-mediated sensitisation to amoxicillin.** A disk diffusion experiment and corresponding plate map demonstrating loss of sensitivity to amoxicillin and relebactam, and meropenem and relebactam in *M. abscessus* pVV16-$Bla_{Mab}$ (S1a). This change in sensitivity can be attributed to the overexpression of $Bla_{Mab}$ as the empty vector control (S1b) exhibited no change in sensitivity from the WT strain.

S2(a)

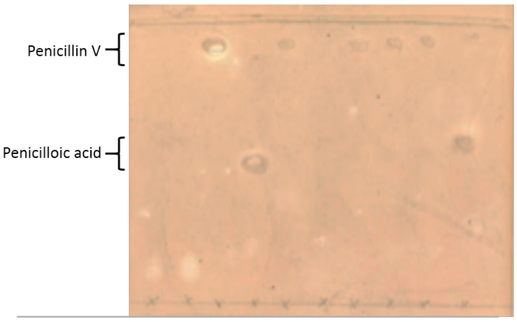


| | 1 | 2 | 3 | 4 | 5 | 6 | 7 | 8 | 9 | 10 |
|---|---|---|---|---|---|---|---|---|---|---|
| Buffer | X | X | | X | X | | X | X | X | X |
| Penicillin V | | | X | X | X | | X | X | X | X |
| $Bla_{Mab}$ | | X | | X | X | | X | X | X | X |
| Avibactam | | | | | X | | | | | |
| Relebactam 0s | | | | | | X | | | | X |
| Relebactam 10s | | | | | | | X | | | |
| Relebactam 30s | | | | | | | | X | | |
| Relebactam 60s | | | | | | | | | X | |

S2(b)

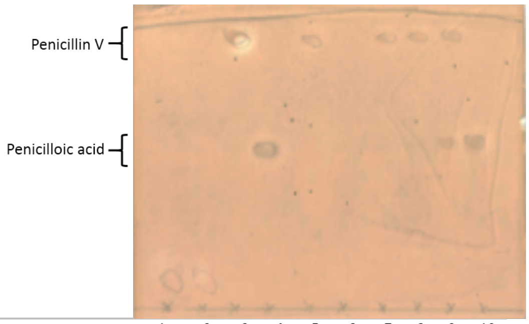


| | 1 | 2 | 3 | 4 | 5 | 6 | 7 | 8 | 9 | 10 |
|---|---|---|---|---|---|---|---|---|---|---|
| Buffer | X | X | | X | X | | X | X | X | X |
| Penicillin V | | | X | X | X | | X | X | X | X |
| $Bla_{Mab}$ | | X | | X | X | | X | X | X | X |
| Avibactam | | | | | X | | | | | |
| Relebactam 200 µg/mL | | | | | | X | X | | | |
| Relebactam 20 µg/mL | | | | | | | | X | | |
| Relebactam 2 µg/mL | | | | | | | | | X | |
| Relebactam 0.2 µg/mL | | | | | | | | | | X |

**Supplementary Figure 2: Our novel Thin Layer Chromatography (TLC) assay exhibiting the activity of $Bla_{Mab}$ in the turnover of penicillin V (high $R_f$ value) to penicilloic acid (lower $R_f$ value).** In the absence, or termination of activity of $Bla_{Mab}$ (by boiling (100 °C for 1 h) or addition of known inhibitor avibactam (Lefebvre et al., 2017) (200 µg/mL) no lower $R_f$ value spot corresponding to penicilloic acid is seen on the TLC plate. The addition of relebactam to the reaction between $Bla_{Mab}$ and penicillin V also results in the absence of the lower Rf value spot, suggesting inhibition of $Bla_{Mab}$. This inhibitory activity was seen within 10 seconds of pre-incubation of relebactam with $Bla_{Mab}$, before addition of penicillin V, in a time course TLC assay. However, the addition of relebactam at the same time (t=0) as penicillin V resulted in a lack of inhibitory activity (S2a). The minimum concentration of relebactam required for inhibition of $Bla_{Mab}$ in the TLC activity assay was assessed using a range of concentrations (200, 20, 2 and 0.2 µg/mL). Activity of $Bla_{Mab}$ was maintained below a relebactam concentration of 2 µg/mL in the TLC activity assay, suggesting a minimal concentration of relebactam required to inhibit $Bla_{Mab}$ in the assay is within the range of 20 to 2 µg/mL (S2b).

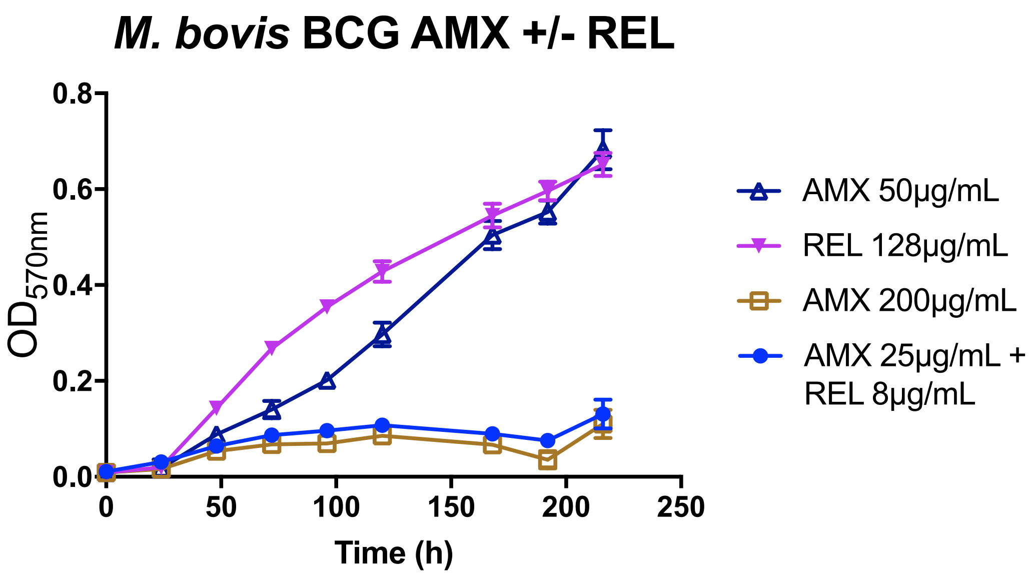


**Supplementary Figure 3: The combination of amoxicillin (AMX) and the β-lactamase inhibitor relebactam (REL) significantly improves activity against *Mycobacterium bovis* BCG.** Growth curves were conducted with *Mycobacterium bovis* BCG Pasteur in medium containing 128 µg/mL REL and 50 µg/mL AMX, neither of which exhibited inhibitory activity. The minimal inhibitory concentration observed for AMX was 200 µg/mL, however with the addition 8 µg/mL REL, the MIC of AMX was reduced to 25 µg/mL. This result suggests potential for this combination in the treatment of tuberculosis as well as *Mycobacterium abscessus* infection.
